# A tissue-aware machine learning framework enhances the mechanistic understanding and genetic diagnosis of Mendelian and rare diseases

**DOI:** 10.1101/2021.02.16.430825

**Authors:** Eyal Simonovsky, Moran Sharon, Maya Ziv, Omry Mauer, Idan Hekselman, Juman Jubran, Ekaterina Vinogradov, Chanan M. Argov, Omer Basha, Lior Kerber, Yuval Yogev, Ayellet V. Segrè, Hae Kyung Im, GTEx Consortium, Ohad Birk, Lior Rokach, Esti Yeger-Lotem

## Abstract

Genetic studies of Mendelian and rare diseases face the critical challenges of identifying pathogenic gene variants and their modes-of-action. Previous efforts rarely utilized the tissue-selective manifestation of these diseases for their elucidation. Here we introduce an interpretable machine learning (ML) platform that utilizes heterogeneous and large-scale tissue-aware datasets of human genes, and rigorously, concurrently and quantitatively assesses hundreds of candidate mechanisms per disease. The resulting tissue-aware ML platform is applicable in gene-specific, tissue-specific, or patient-specific modes. Application of the platform to selected Mendelian disease genes pinpointed mechanisms that lead to tissue-specific disease manifestation. When applied jointly to diseases that manifest in the same tissue, the models revealed common known and previously underappreciated factors that underlie tissue-selective disease manifestation. Lastly, we harnessed our ML platform toward genetic diagnosis of tissue-selective rare diseases. Patient-specific models of candidate disease-causing genes from 50 patients successfully prioritized the pathogenic gene in 86% of the cases, implying that the tissue-selectivity of rare diseases aids in filtering out unlikely candidate genes. Thus, interpretable tissue-aware ML models can boost mechanistic understanding and genetic diagnosis of tissue-selective heritable diseases. A webserver supporting gene prioritization is available at https://netbio.bgu.ac.il/trace/.

## INTRODUCTION

Among the major challenges in genetic study of Mendelian and rare heritable diseases are identification of pathogenic variants^1^ and the molecular mechanisms by which they lead to disease phenotypes^2^. Exome and whole-genome sequencing of patients typically yield large numbers of candidate variants of unknown significance (VUS)^3^. These VUS are scrutinized via multiple strategies, including sequence and conservation analyses (e.g.,^4–7^), comparison to exome/genome sequencing databases or known pathogenic variants^8–10^, and phenotypic resemblance to known disease-associated genes^11,12^ or phenotypes^13^. However, the rate of successful genetic diagnostic of patients stands on 25-60%^14^, calling for additional strategies.

When a pathogenic variant is identified, the molecular mechanisms by which it leads to disease phenotypes typically remain elusive. Attesting to the complexity of this challenge, molecular mechanisms have remained hidden even for Mendelian diseases with long known and well-established pathogenic variants^15–19^. Especially intriguing are mechanisms that underlie tissue-selective Mendelian diseases^2^. These diseases manifest clinically in few tissues, whereas the gene harboring the pathogenic variant is often expressed broadly across the human body^2^.

Recent years were marked by immense molecular characterizations of tens of physiological human tissues, including transcriptomes^20^, proteomes^21^, and regulatory and epigenetic signals^22^. These datasets were harnessed to confront the above challenges. Genes harboring pathogenic variants (denoted disease genes) were shown to be preferentially expressed^23,24^ or to participate in molecular interactions that occurred preferentially^24–28^ in their disease-susceptible tissues. Likewise, genes associated with complex traits were shown to disturb regulatory relationships^29^, gene modules^30^, or to overlap with eQTLs^31–33^ that were active in trait-manifesting tissues. Additionally, omics datasets were used in tools for prioritizing candidate disease genes based on their resemblance to seed genes associated with disease phenotypes (Endeavor^11^, pBRIT^12^, GADO^13^). Recently, annotation of exon expression across tissues was shown to improve interpretation and prioritization of rare variants^9,10^. Yet, a framework that simultaneously assesses numerous molecular mechanisms, and supports the inference of disease mechanisms and disease genes has been lacking.

Machine learning (ML) offers a framework for simultaneous exploration of heterogeneous datasets (reviewed in ^34^). For example, ML helped identify interactome-rewired genes in tissue-specific cancers^27^, infer gene modules for complex diseases^35^, and predict diabetes-causing variants from epigenomes of pancreatic islets^36^. Here we present an interpretable ML framework, termed ‘Tissue Risk Assessment of Causality by Expression’ (TRACE), for improving molecular understanding of tissue-selective disease manifestation and genetic diagnosis of patients. The framework was based on 4,744 tissue-aware gene features that were deduced from physiological and heterogeneous datasets and was tested by using 1,105 disease genes that lead to tissue-specific diseases. Gene-specific models identified mechanisms that were indeed associated with the tissue-specific manifestation of the respective diseases. Tissue-specific models that jointly analyzed multiple diseases pointed not only to known tissue-specificity mechanisms, but also revealed the dominant contribution of tissue-selective cellular processes. Lastly, we applied TRACE to prioritize candidate VUS-containing genes identified in patients with rare tissue-specific Mendelian diseases. Despite being oblivious to the actual genetic variation, in 86% of the cases TRACE successfully prioritized the verified pathogenic gene. Thus, TRACE can complement and enhance existing VUS-interpretation tools. A TRACE webserver for prioritizing candidate disease-associated genes based on cross-validation assays is available at https://netbio.bgu.ac.il/trace/.

## RESULTS

### Constructing tissue-aware features and dataset for ML

We constructed a large-scale dataset consisting of 4,744 engineered (interpretable) and abstract (performance boosting) tissue-aware features per protein-coding gene (Fig. 1A, Table S1, Dataset S1). Tissue-aware features were derived from diverse data sources, predominantly transcriptomes of mature^20^ and developing^37^ physiological human tissues, as well as tissue eQTLs^20^, experimentally-detected physical protein-protein interactions (PPIs) (e.g.,^38,39^), and Gene Ontology (GO) biological process terms and annotations^40^. Features were inferred from a single data source or by integrating multiple data sources. For example, we inferred tissue PPIs^24^ by integrating tissue transcriptomes and PPIs. In addition to including features with known association to tissue-selectivity, we introduced novel features. For example, we quantified the activity of cellular processes per tissue by integrating GO terms with the tissue expression levels of GO term-associated genes (Methods). To support interpretability, we engineered (computed) tissue-comparative features, such as tissue-preferential gene expression^41^, tissue-differential PPIs^28^, and tissue-differential process activity (Methods). To enhance the predictive power of ML models by reducing interactomics data-loss we added abstract network embedding vectors that represented gene neighborhoods in tissue interactomes (Methods). Missing values were imputed and values were transformed and scaled (Methods).

**Fig. 1.**
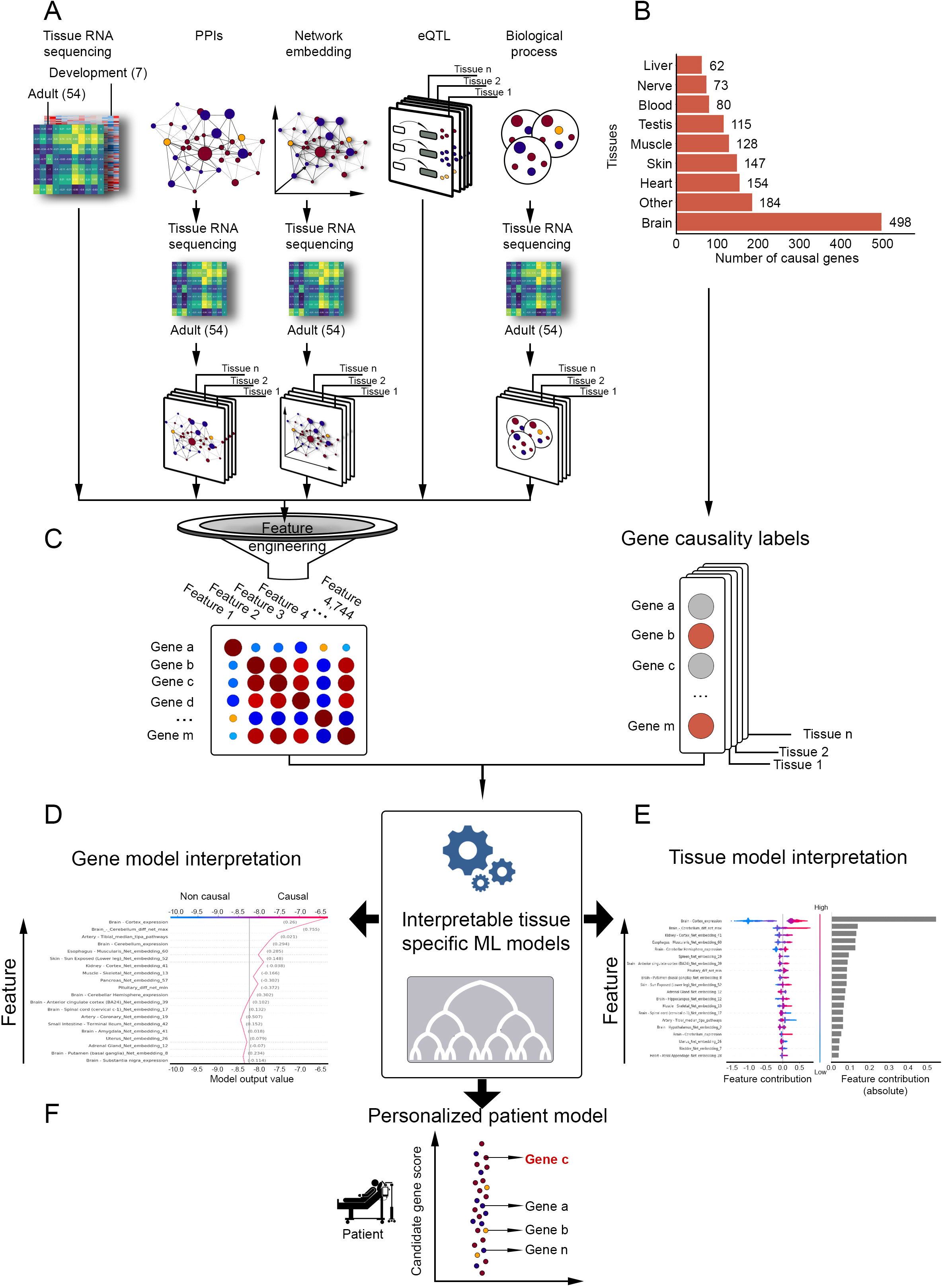
The ML scheme for interpreting tissue-selective disease mechanisms. A. Construction of the tissue-aware gene features dataset used in the analysis. Starting with datasets including transcriptomes of 54 human adult tissues and seven developing organs, PPIs, tissue eQTLs and gene annotations to biological processes, datasets were integrated and interpretable features were engineered, resulting in 4,744 features for 18,927 protein-coding genes. B. The number of protein-coding genes known to be causal for Mendelian diseases that manifest in the designated tissues or in other tissues (marked ‘Other’). C. The input to interpretable decision-trees tissue models includes the dataset of tissue-aware gene features and gene-causality labels for the modeled tissue. Red labels mark genes causal for Mendelian diseases that manifest in the modeled tissue. Various outputs of the models are described in panels D-F. D. Buildup of the output value for the predicted causality of a query gene in the modeled tissue. Starting with a neutral value at the bottom, the output value accumulates according to the feature values of the query gene. Features are ordered from bottom to top by their increased contribution to the model, allowing interpretation. The location of final output value on the X axis indicates whether the query gene is predicted to be causal (right) or not causal (left) for a disease that manifests in the modeled tissue (the gray vertical bar denotes the baseline value). E. A quantitative view of the contribution of features to a tissue model. Features are ordered from bottom to top by their increased absolute contribution to the model (grey bars), allowing interpretation. Per feature, each dot represents the feature value of a different gene. Dots are colored red for high values and blue for low values, and spread horizontally from left to right by their causality for a disease that manifests in the modeled tissue. F. Given a list of candidate disease-causing genes of a patient as input, the model outputs scores representing the predicted probabilities of the candidate genes to cause a disease that manifests in the patient`s affected tissue (gene C represents the verified causal gene).

Next, we labeled genes according to their causality for tissue-selective Mendelian disorders. We retrieved Mendelian disease genes from OMIM, and combined them with manually curated associations between Mendelian diseases and clinically-affected tissues^28,42^. Altogether, our features dataset covered 18,927 protein-coding genes, including 1,105 curated disease genes that unitedly affected 22 tissues (Table S2). Eight of the affected tissues were each associated with over 60 disease genes, including blood, brain, heart, liver, nerve, skeletal muscle, skin and testis (Fig. 1B), summing up to 1,031 disease genes henceforth referred to as tissue-casual. For each of the eight tissues, we labeled genes to denote whether they lead to a disease that clinically affects that tissue. Other tissues had fewer than the required number of disease genes for training the ML models (<60 genes). The tissue-aware dataset and gene causality labels provided the basis for ML (Fig. 1C), which we applied to create gene-specific, tissue-specific, and personalized patient models (Fig. 1D-F).

### Gene-specific ML models illuminate disease mechanisms

Our first goal was to identify features that contribute to tissue-selective manifestation of a certain disease. We used the XGBoost^43^ (XGB) gradient boosting method (Fig. 1C), since decision forest methods perform well on tabular data with thousands of training instances^44^, imbalanced classification tasks^45^ and high-dimensional data that include large numbers of dependent features^46^. We then applied XGB to illuminate the tissue-specific impacts of disease genes. Specifically, for each disease gene we created a distinct XGB model by training the model on all other genes, which we labeled according to their causality in the respective disease tissue. We then applied the trained model to the query disease gene (Methods). To rigorously highlight the features that contributed to the decision of each model, we used the SHAP (SHapley Additive exPlanations) algorithm^47^, which is a game theoretic method for explaining the prediction of ML models.

We modeled four genes that lead to diseases manifesting in brain, skin, heart and skeletal muscle (Fig. 2). Each model successfully classified the query disease gene as causal in its respective disease tissue. The model for the epilepsy gene SYN1, which we trained using brain labeling, pointed to ‘cerebellum differential PPIs’ and ‘brain cortex expression’ (Fig. 2A), in accordance with the exceptionally high expression level of SYN1 in brain versus other tissues (Fig. 2A). The model for the psoriasis gene IL36RN, which we trained using skin labeling, pointed to ‘skin expression’ as the main contributing feature (Fig. 2B), in accordance with IL36RN exceptionally high expression level in skin versus other tissues (Fig. 2B). Especially revealing were the models for CACNA1C that leads to arrhythmia and DMD that leads to Duchenne muscular dystrophy. These genes were not expressed exceptionally high in their respective disease tissues. Instead, the top contributing feature of each gene model was the differential process activity of the gene within its disease tissue, which was high relative to other tissues (Fig. 2C,D). Examination of the processes that contributed to these features revealed that they were upregulated specifically in the disease tissue. In case of CACNA1C, these processes were related to electrical conduction of the heart which is perturbed in arrhythmia, such as ‘membrane depolarization during atrial cardiac muscle cell action potential’ (Fig. 2C). In case of DMD, the process was ‘muscle filament sliding’, which was indeed shown to be impaired in mdx mouse model for Duchenne^48^. These examples demonstrate that gene-specific models can point to gene-centered mechanisms (Fig. 2A,B) or to mechanisms involving a group of genes whose combined function underlies disease phenotypes (Fig. 2C,D).

**Fig. 2.**
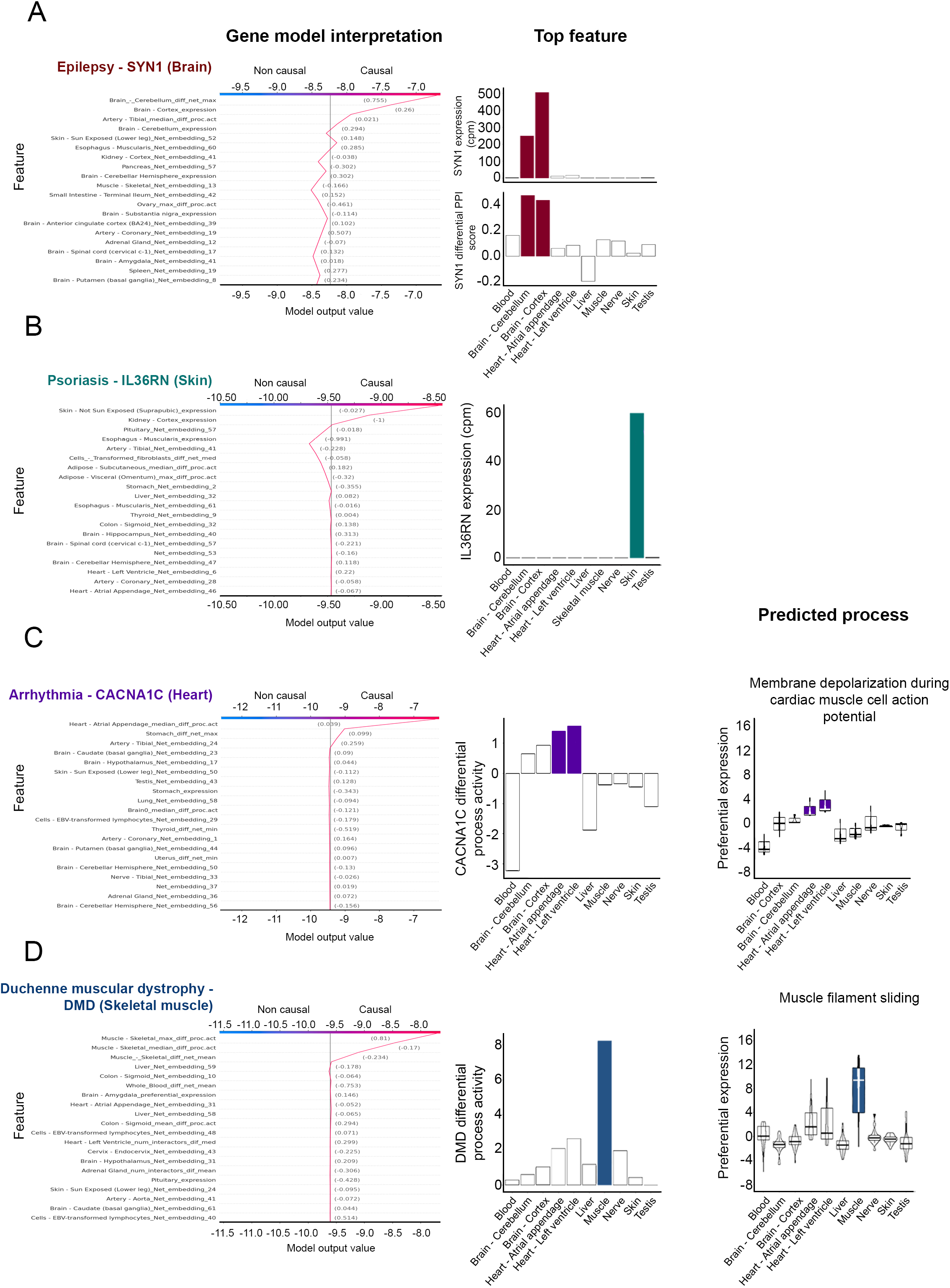
Gene-specific models illuminate gene causality mechanisms. A. The model for the epilepsy gene SYN1 predicted its causality in brain (left). The model’s topmost contributing features, ‘brain cortex expression’ and ‘cerebellum differential PPIs’, agree with the exceptionally high expression level and differential PPIs score of SYN1 in brain cortex and cerebellum relative to other tissues (middle, purple bars). B. The model for the psoriasis gene IL36RN predicted its causality in skin (left). The model’s topmost contributing feature, ‘skin expression’, agrees with IL36RN exceptionally high expression level in skin relative to other tissues (right, green bar). C. The model for the arrhythmia gene CACNA1C predicted its causality in heart (left). The model’s topmost contributing feature, ‘differential process activity in heart’, fits with the high differential process activity of CACNA1C in heart relative to other tissues (middle). This high activity stems from the preferential expression in heart of genes composing arrhythmia–related processes such as ‘membrane depolarization during atrial cardiac muscle cell action potential’ (right). D. The model for the Duchenne muscular dystrophy gene DMD predicted its causality in skeletal muscle (left). The model’s topmost contributing feature, ‘differential process activity in skeletal muscle’, agrees with the exceptionally high differential process activity of DMD in skeletal muscle relative to other tissues (middle). The high activity was due to the preferential expression in skeletal muscle of genes composing the process ‘muscle filament sliding’ (right), previously implicated in Duchenne.

### Tissue-specific ML models reveal dominant tissue-selectivity mechanisms

Encouraged by the interpretability of gene-specific models, we extended our interpretability analysis to identify features that commonly contributed to the tissue-selectivity of multiple genes. We therefore trained a separate XGB classification model for each of the eight tissues with over 60 genes (Fig. 1B) that aimed to distinguish tissue-casual genes from other genes (Methods). We then assessed the performance of each model by using 10-fold cross-validation. The average area under the receiver operating characteristic curve (AUC) obtained by the various tissue models was 0.71-0.87 (Fig. S1), attesting to their discriminative power. Next, we used SHAP to assess the contribution of each feature to each tissue model (Fig. 3A, Fig. S2). Features that were associated with the modeled tissue, e.g., ‘brain cortex expression’ in the brain model and ‘differential process activity in skin’ in the skin model, were among the ten most important features per model, and among the top three most important features in 6/8 models. These features were complemented by features of other tissues. For example, the third most contributing feature in the skin model was ‘differential process activity in subcutaneous fat’, a tissue located just beneath the skin (Fig. 3A). Interestingly, the prediction of skin-causal genes was associated with high gene values for the skin-associated feature and low gene values for the feature associated with subcutaneous fat.

**Fig. 3.**
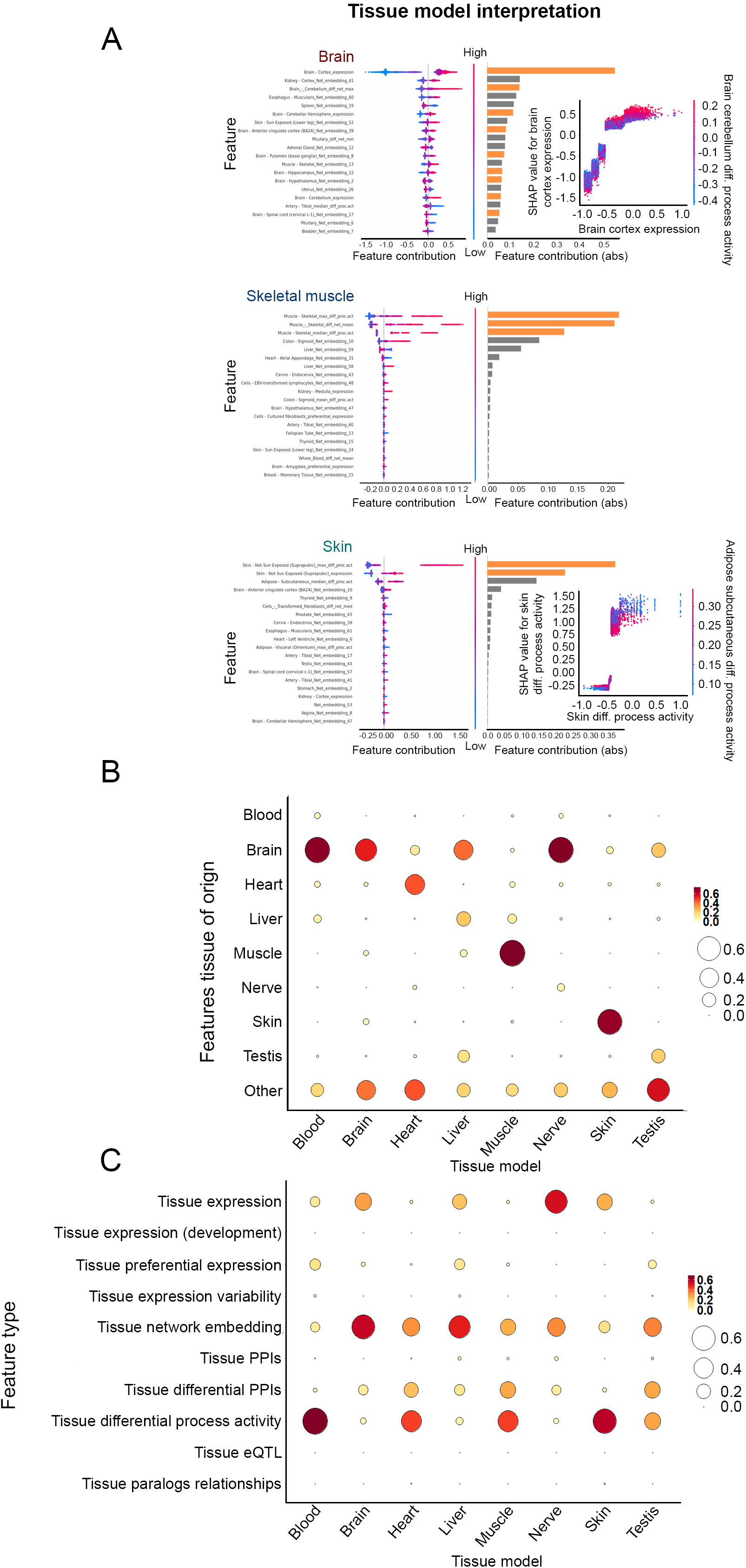
Tissue-specific models reveal common tissue-selectivity mechanisms. A. The 20 topmost contributing features to the brain, skin and skeletal muscle models (see legend of Fig. 1E). Features of the modeled tissue are marked by orange bars and are among the topmost features. Inset panels in the brain (top) and skin (bottom) models show the topmost contributing feature to the model (X and Y axes) and the feature that best complemented it (right-most axis). Dots represent gene values and contribution for the topmost feature and are colored by gene values on the complementary feature. Detailed tissue models appear in Fig. S2. B. The contribution of tissue-aware features to tissue models upon aggregating features by their associated tissue. Sub-regions of a tissue were associated with their main tissue. Aggregated features of the modeled tissue were among the three topmost contributing features in the brain, heart, liver, skeletal muscle, skin and testis models. C. The contribution of features to tissue models upon aggregating features by their type. Tissue differential process activity, tissue network embedding, and tissue expression were the topmost contributing feature types in at least one model.

Next, we used the rigorous quantification by SHAP of the contribution to tissue-selectivity of the thousands of features in our dataset to identify common determinants of tissue-selectivity in an unbiased manner. For this, we summarized the contribution of the different features per tissue model by aggregating their normalized SHAP importance (Methods). We first summarized the contribution of features that were associated with the same tissue, e.g., ‘brain cortex expression’ was associated with brain (Fig. 3B). In 7/8 tissue models, the modeled tissue itself was among the top three out of 54 tissues with the highest contribution to its model, implying that the tissue-causality of a gene frequently stems from the gene’s behavior in its disease-susceptible tissue.

To further obtain a mechanistic insight into common determinants of tissue-selectivity, we summarized the contribution of features of the same type per tissue model, regardless of their associated tissue. For example, ‘brain cortex expression’ and ‘liver expression’ were associated with the mechanism ‘expression’ (Fig. 3C). Expression level, indeed a major and well-established determinant of tissue-selectivity^8^, was the topmost mechanism in one model. Network neighborhood, another recognized determinant of tissue-selectivity, was topmost in 3/8 models. Notably, differential process activity, which was not previously recognized as a driver of tissue-selectivity, was the topmost mechanism in 4/8 tissue models, attesting to its wide relevance. This implies that, in many cases, the pathogenic gene is not the sole driver of disease, but rather it is part of a process whose integrity is essential for tissue physiology. Moreover, identification of this process provides insight into the actual disease mechanisms and thus could open avenues for therapy (Fig. 2C,D). Differential PPIs and preferential expression were also influential across tissues, suggesting that both absolute (e.g., expression) and relative (e.g., preferential expression) tissue-aware gene features affect the tissue-selectivity of diseases.

### Tissue-specific ML models prioritize genes underlying tissue-selective diseases

We turned to harness our ML framework toward prioritization of candidate tissue-causal disease genes in each of the eight disease-susceptible tissues. For this, we created a robust two-layer ML scheme, denoted ‘Tissue Risk Assessment of Causality by Expression’ (TRACE). The first layer employed five ML classifiers that we found to perform complementary well: logistic regression (LR), a multilayer perceptron (MLP) neural network, and three tree-based ensemble methods that included XGB, random forest (RF) and gradient boosted trees initiated by a logistic regression model (LR+GB). Per classifier, we applied 10-fold cross-validation to compute the probability of each gene to be tissue-causal, and then scaled the probabilities to values between 0-10 (Fig. 4A, Methods). In the second layer, we employed a deep neural network meta-learner^49^ (meta-MLP). The meta-MLP received as input the gene values obtained by the five classifiers, and produced a final TRACE score, also scaled between 0 and 10.

**Fig. 4.**
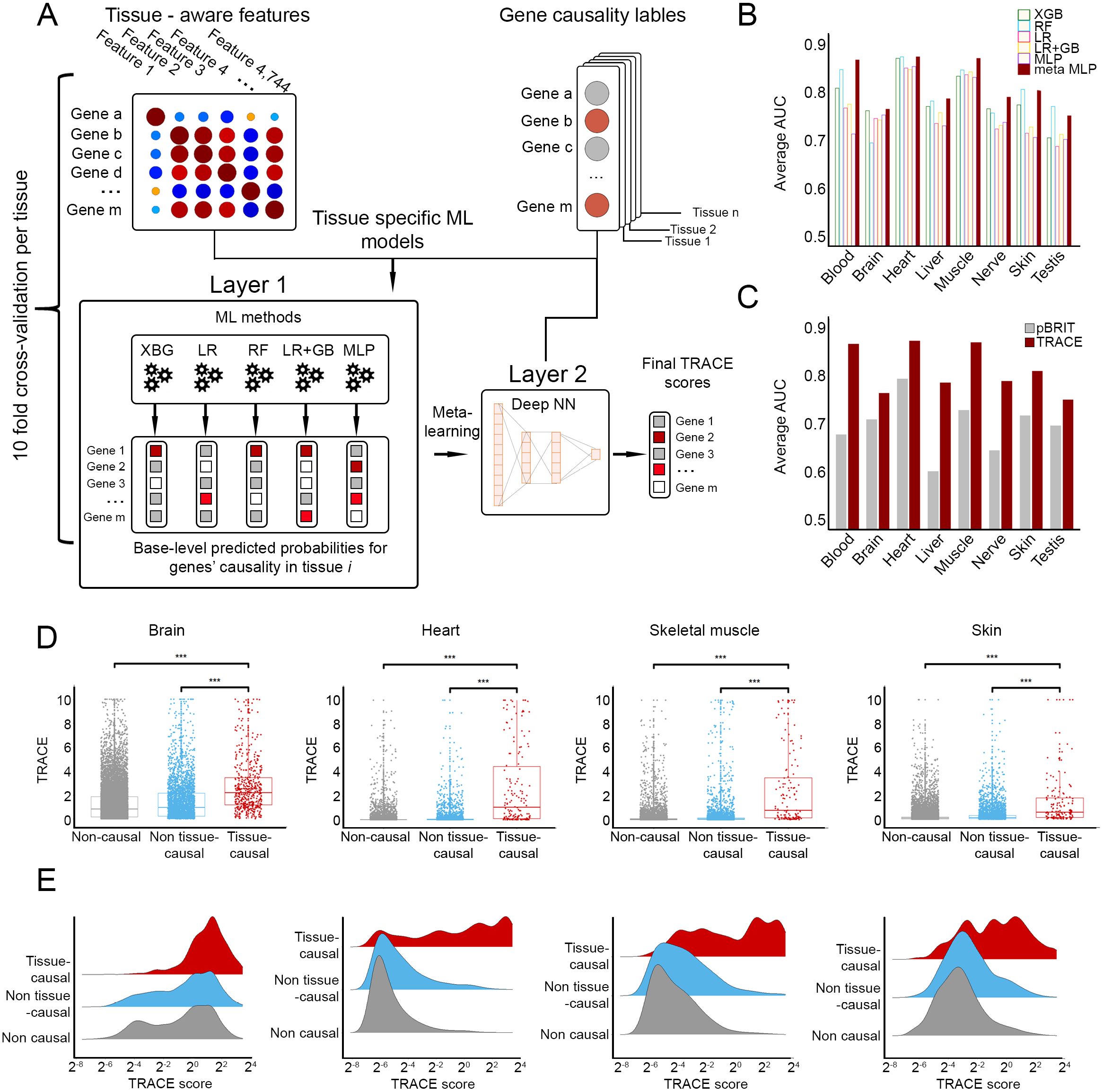
TRACE prioritization of genes associated with diseases that manifest in distinct tissues. A. A schematic view of TRACE. Input to TRACE included the dataset of tissue-aware gene features and gene-causality labels for the modeled tissue (red labels mark genes causal for Mendelian diseases that manifest in the modeled tissue). The first layer includes independent application of five ML methods. Each method outputs gene scores reflecting the predicted probability that the gene causes a disease that manifests in the modeled tissue. The second layer uses gene scores as input to a neural network (NN) meta-learner that produces the final TRACE score (red labels mark genes with higher predicted causality scores). B. The average AUC obtained per ML method and tissue model. The highest AUC were typically obtained by the meta-learner. C. The average AUC obtained with pBRIT (grey) and TRACE (crimson) per tissue model. D. Gene TRACE scores in brain, heart, skeletal muscle and skin models. Each dot represents a different gene. Genes were divided into non-causal genes (grey), genes that are causal but not in the modeled tissue (non tissue-causal, blue), and genes causal in the modeled tissue (tissue-causal, red). Tissue-causal genes had significantly higher TRACE scores compared to non-causal genes and to tissue non-causal genes (***=3E-16; MW, adjusted p-values). E. Density ridge plots of gene TRACE scores in brain, heart, skeletal muscle and skin models. Tissue-causal genes (red) were over-represented among genes with high TRACE scores. Plots for additional tissue models appear in Fig. S5.

Classifiers’ performance was similar between ML methods and across tissues (Fig. S3), with the meta-MLP predominantly producing the top AUC (0.75-0.87, Fig. 4B) and top precision-recall scores (Fig. S4A). We compared the final scores of TRACE to pBRIT, a recent tool for variant prioritization that applies intermediate integration data fusion^12^ (Methods). Despite pBRIT relying on more types of functional and phenotypic gene annotations, TRACE performance was favorable, especially with respect to AUC (Fig. 4C, Fig. S4B). To further assess the tissue-specificity of TRACE, we divided genes into three separate groups per tissue: (i) tissue-causal genes, (ii) causal genes that were not associated with the susceptible tissue (denoted tissue non-causal), and (iii) non-causal genes (denoted non-causal). We then compared between the TRACE scores of the different groups (Fig. 4D, Fig. 4E, Fig. S5). Across models, tissue-causal genes were enriched among genes with high TRACE scores. Tissue-causal genes not only had significantly higher TRACE scores when compared to non-causal genes, but also when compared to tissue non-causal genes (p≤4.4E-13 and p≤1.1E-6, respectively, Mann-Whitney U test (MW)). The ability of the models to favor tissue-causal genes over genes that are causal for disease that manifest in other tissues attests to the tissue-specificity of the models.

So far, we applied TRACE to susceptible tissues that were physiologically distinct from each other. Next, we turned to analyze the complex and often-uncertain selectivity of brain diseases to brain regions^50^. We divided brain into six regions with available transcriptomic profiles, including cortex, cerebellum, basal ganglia, spinal cord, hypothalamus, and amygdala (Table S3). We manually associated brain diseases to susceptible brain regions based on anatomical findings (see Methods). Altogether, we associated with medium to high confidence 594 diseases and 532 causal genes to susceptible brain regions, of which only cortex and cerebellum harbored over 60 genes each and were henceforth modeled (Fig. S6A, Table S3). As observed for physiologically distinct tissues, TRACE models were discriminative (AUC of 0.77 in both). Likewise, cortex-causal and cerebellum-causal genes had significantly high TRACE scores when compared to non-causal and non-brain causal genes (p≤3.3E-16, MW, Fig. S6B,C). Notably, the TRACE scores of cortex-causal and cerebellum-causal genes remained significantly high relative to brain-causal genes that were not cortex-causal or cerebellum-causal, respectively (adjusted p-value of 0.028 and 0.033, respectively, MW, Fig. S6B). Some of the high-scoring genes were related with the appropriate brain region, however with low confidence. For example, GNB1, a gene causal for mental retardation, and GAN, a gene causal for giant axonal neuropathy, had a low confidence association with brain cortex, where they indeed ranked high (TRACE scores of 8.62, top 0.2%, and 10, respectively). Thus, TRACE models favor tissue-causal genes over genes that are causal for disease that manifest in physiologically-related tissues.

To facilitate similar analyses we developed the TRACE webserver (http://netbio.bgu.ac.il/trace/). Users can upload a gene list or a VCF file and select a disease-susceptible tissue, and obtain TRACE scores for the respective query genes in the user-selected tissue, allowing their prioritization.

### Patient-specific ML models can boost genetic diagnosis of rare-disease patients

To test whether TRACE could be beneficial for genetic diagnosis of patients with rare tissue-specific Mendelian diseases, we applied TRACE to test data from 50 diagnosed patients. Each patient underwent whole-exome or whole-genome sequencing and VUS characterization as done routinely in clinical genetic diagnosis or discovery (Fig. 5A). Per patient, we compiled a list of VUS-containing genes, referred to as query genes. Next, we created a TRACE model per patient and disease-affected tissue (five patients had two affected tissues), which we trained on all genes except for the patient’s query genes. We then applied the trained model to the patient’s query genes to predict their likelihood to be causal in the disease-affected tissue, and prioritized them by their TRACE scores (Fig. 5B).

**Fig. 5.**
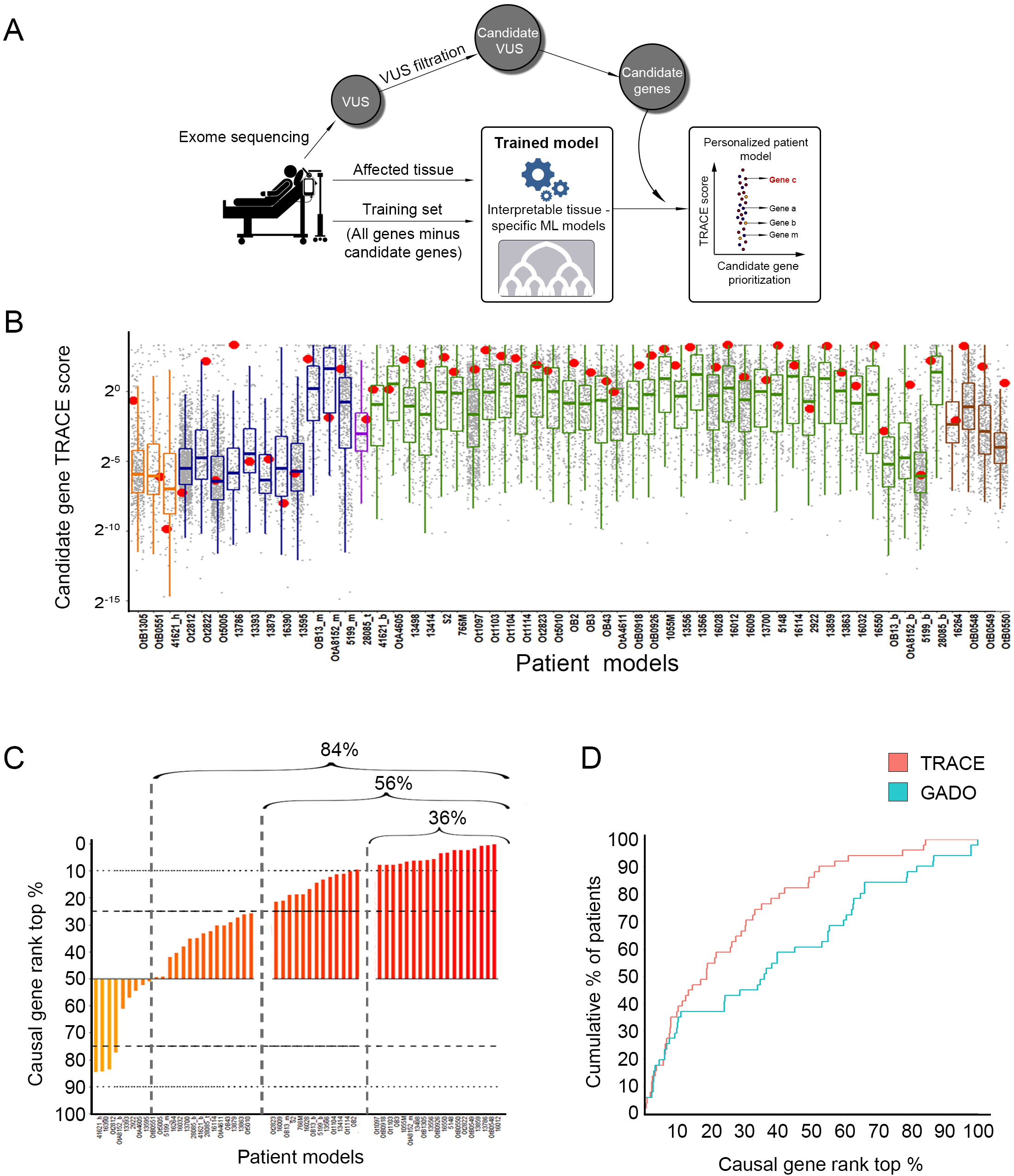
TRACE prioritization of patient-derived candidate genes associated with rare diseases. A. A schematic view of a patient-tailored TRACE modeling. VUS identified via exome sequencing of the patient were filtered by standard approaches, and remaining VUS were mapped to their harboring genes, denoted candidate genes. Next, A TRACE model was trained on all genes except for the patient’s candidate genes, using gene labels that match the affected tissue of the patient. Lastly, the trained model was applied to the patient’s candidate genes, which were then prioritized according to their TRACE scores. The red-labeled gene contains the patient’s pathogenic variant. B. The TRACE scores of candidate genes in 55 models for 50 patients with tissue-specific rare diseases (five patients were modeled in two distinct tissues). Per model, each dot represents a candidate gene; the red dot marks the verified pathogenic gene. Boxplot color reflect the modeled tissue (orange=heart, blue=skeletal muscle, purple=testis, green=brain, brown=skin). In 46/55 models the disease-causing gene was ranked above the median. C. The rank of the verified pathogenic genes out of the patient’s candidate genes. Per model, ranks were determined based on gene TRACE scores, such that top scoring gene was ranked first. 84% of the verified disease genes ranked above the median; 56% of the verified disease genes ranked at the top quartile; and 36% of the verified disease genes ranked at the top 10%. D. Comparison between the rank of the verified pathogenic genes out of the patient’s candidate genes between TRACE and GADO^13^. TRACE prioritizations were significantly better (median rank of 36 versus 59, p=0.0085, paired Wilcoxon).

Though TRACE analysis of query genes was literature blind and relied only of large-scale profiling data, several well-established disease genes (some containing novel pathogenic variants) ranked at the top 1%. Examples include OPA1 that was verified in a patient with optic atrophy and neuropsychiatric disorders (MIM #125250); Sarcoglycan Gamma (SGCG) that was verified in a patient with muscular dystrophy (MIM #253700); and tumor protein p63 (TP63) that was causal for ectodermal dysplasia syndrome (MIM #103285, #604292).

Several newly discovered disease genes, causing novel phenotypes in the queried populations, demonstrate the utility of TRACE. In one example, a familial case of lethal, severe microcephaly with various neurological features, was found to have a unique mutation in SEC31A, yet to be described as causing human disease^51^. TRACE analysis of data from this patient ranked SEC31A as the eighth most likely causal gene out of 107 candidate genes, mostly owing to its high expression in brain cortex. In a different case, a familial syndrome of muscle hypotonia, failure to thrive, and developmental delay, a variant in PAX7 was recently identified as causing the disease^52^. This disease affects multiple organs but manifests most severely in skeletal muscle. TRACE ranked PAX7 in skeletal muscle TRACE as the 10th most likely causal gene out of 150 candidates. These examples demonstrate the utility of TRACE in the complex task of clinical prioritization and preliminary interpretation of disease-associated variants.

Analysis of TRACE across all patients revealed the boosting capability of TRACE. The verified disease gene (i.e., the gene containing the verified pathogenic variant) ranked above the median in 84% of the cases, implying that low ranking candidate genes can typically be ignored (Fig. 5C). In 56% of the cases, the verified disease gene ranked at the top quartile, and in 36% of the cases it ranked at the top 10%. Verified disease genes ranked especially high in models of patients with brain and skin diseases (64% and 75% ranked at the top quartile, respectively). Lastly, we compared TRACE prioritization to the prioritization offered by GADO, a recently published tool that uses human tissue transcriptomes to prioritize genes causing specific disease phenotypes ^13^. TRACE prioritization was significantly better than that of GADO (Fig. 5D, p=0.0085, paired Wilcoxon), especially in prioritizing more elusive genes.

## DISCUSSION

We presented a tissue-aware ML framework for prioritizing candidate disease genes and the mechanisms by which they lead to disease. For this, we integrated heterogeneous omics datasets, creating engineered features that were motivated by previous studies of tissue-selectivity^2,23–26,29–31,42,53^, and abstract interactomics features (Table S1). By employing early integration, rather than late^11^ or intermediate^12^ integration techniques, the resulting gene (Fig. 2) and tissue models (Fig. 3A) enabled feature-based interpretation, and uncovered complementary features that originated from different datasets^54^.

TRACE gene-specific models were able to provide mechanistic insight into the tissue-selective impact of genes (Fig. 2). Moreover, the dominant features revealed by TRACE in tissue-specific models generally agreed with known tissue selectivity factors, such as gene expression levels and interactome neighborhoods in disease-susceptible tissues^23,24^ (Fig. 3B,C). Yet, some previously acknowledged factors, such as eQTLs^31,53^ or paralog levels^42^, were not highlighted. These features might have been masked by other features, or their signal could be low in the heritable diseases dataset that we used^55^. Notably, TRACE exposed the differential activity of biological processes as a dominant tissue-selectivity feature that was not highlighted before (Fig. 3C). This feature provided meaningful insight into the specific processes actually perturbed in disease (Fig. 2C,D). Together with preferential expression and differential PPIs, these tissue-comparative features support a holistic examination of ubiquitously-expressed causal genes beyond the context of susceptible tissues.

When applied to prioritize disease-causing genes per tissue, TRACE models were tissue-sensitive (Fig. 4) even when applied to distinct brain regions (Fig. S6). For the brain regions analysis, we manually curated 532 brain diseases to affected brain sub-regions, creating a valuable resource for further analyses (Table S3). An online TRACE webserver for gene prioritization based on cross validation assays is available at http://netbio.bgu.ac.il/trace/.

In contrast to common VUS prioritization schemes^1^, TRACE application to data from patients did not include variant information. Nevertheless, TRACE ranked the pathogenic gene high in most patient models (Fig. 5), attesting to the importance of considering the affected tissue of the patient when analyzing VUS. Since common VUS prioritization schemes are almost oblivious to tissue contexts, TRACE provides a powerful complementary addition to current pipelines.

The ML framework that we described could be extended to employ additional ML methods and richer datasets of features, traits and diseases. The limited mechanistic understanding of diseases and the numbers of undiagnosed patients worldwide call for additional ready-to-use tools for boosting disease understanding and diagnosis.

## METHODS

### Gene expression datasets and processing

Transcriptomic profiles of adult human tissues measured via RNA-sequencing were obtained from GTEx v8 and consisted of 17,382 profiles sampled from 54 tissues^20^. Expression values were available for 43,025 genes, including 18,927 protein-coding genes. Raw reads were normalized to obtain the same library size for every sample by using the trimmed mean of M-values (TMM) method by the edgeR package^56^. Genes with at most 10 raw counts in every sample were removed before normalization, as these genes were typically regarded as noise. The normalized expression of a gene in a tissue was set to its median normalized count in the corresponding tissue samples. Genes with median normalized counts ≥ 7 in a tissue were henceforth considered as expressed in that tissue. Only protein-coding genes that were expressed in at least one tissue were further considered.

Preferential expression values per gene and tissue were computed from normalized counts as in Sonawane et al^41^ (see Equation 1). Similarly to Sonawane et al^41^, genes with preferential expression ≥ 2 in a tissue were considered as preferentially expressed in that tissue. Transcriptomic profiles of developing human organs consisted of gene normalized counts that were measured at several time points during development and in adulthood in seven human organs, including cerebrum, cerebellum, heart, kidney, liver, ovary and testis^37^. The expression of a gene in an organ and a time point was set to its median normalized count in samples from the same time point and organ, resulting in a total of 133 profiles.

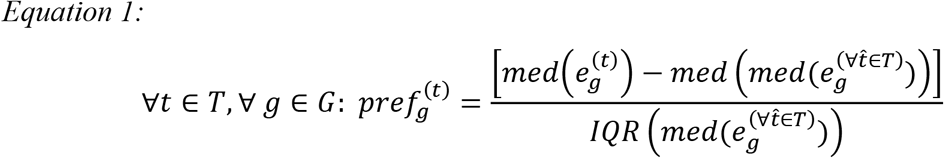

*T* denotes the set of tissues; *G* denotes the set of genes; *pref* denotes preferential expression; *e* denotes normalized count; *IQR* denotes inter-quartile range. For any tissue *t*, 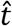 is any other tissue in *T*.

### Construction of the dataset of tissue-aware gene features

The features dataset consisted of 4,744 features per protein-coding gene. Below we describe the different types of features (see Table S1), as well as imputation and scaling of the dataset.

#### Transcriptomic features

Per tissue, each gene was associated with a feature corresponding to its normalized expression in that tissue and a feature corresponding to its preferential expression in that tissue according the GTEx dataset. Per organ that was profiled during development, each gene was associated with features corresponding to its normalized expression in that organ per time point.

#### eQTL features

Data of tissue eGenes, i.e., genes involved in a tissue eQTL, were downloaded from the GTEX portal (March 20th, 2019). Each gene was associated with a feature per tissue that corresponded to its eGene p-value in that tissue.

#### PPI features

Data of experimentally-detected PPIs were downloaded from publicly available databases using the default option of the MyProteinNet web-tool^57^. Each gene was associated with the set of interactors of its corresponding protein. Next, we integrated PPIs with GTEx expression data. Specifically, per tissue, we assigned each gene with three tissue-aware primary PPI-related features, as follows: (i) Tissue interactors: Every gene that was considered to be expressed in the given tissue (i.e., median normalized counts ≥ 7 in that tissue) was assigned with the number of its interactors that were also considered to be expressed in that tissue; otherwise its value was set to 0. (ii) Preferential tissue interactors: Every gene that was considered to be expressed in the given tissue was assigned with the number of its interactors that were considered to be preferentially expressed in that tissue; otherwise its value was set to 0. (iii) Tissue-specific interactors: For every gene with non-zero tissue interactors in that tissue, the gene was assigned with the number of tissue interactors with which it interacted in at most 20% of the tissues (i.e., both the gene and its interactor were expressed in the same tissue in at most 20% of the tissues). For each primary feature (i)-(iii), we calculated two additional features per gene and tissue, to reflect the difference between the primary gene value in that tissue and its expected value. The expected value was calculated in two different ways, once according to the gene’s mean primary value across tissues, and once according to the gene’s median primary value across tissues.

#### Tissue-differential PPI features

Data of differential PPIs per gene and tissue were downloaded from The DifferentialNet database^58^. This database assigns each PPI with a score per tissue that reflects whether the two interacting proteins are over-expressed or under-expressed in that tissue relative to all other tissues. Using these data, we associated each gene with the set of its differential interaction scores per tissue, and created four gene features per tissue, as follows: (i) the minimum differential interaction score of the gene in the given tissue. (ii) The maximum differential interaction score of the gene in the given tissue. (iii) The median differential interaction score of the gene in the given tissue. (iv) The mean differential interaction score of the gene in the given tissue.

#### Network embedding features

Network embedding features were designed to represent the interactome neighborhood of a gene in a given PPI network (interactome). We started by creating a general interactome that contained all experimentally-detected PPIs (see PPI features section above). Next, we integrated the general interactome with GTEx expression data to create tissue interactomes. Specifically, the interactome of a given tissue was set to include all PPIs between protein products of genes that were considered as expressed in that tissue (i.e., genes with median normalized counts ≥ 7 in that tissue). Lastly, we applied network embedding to the general human interactome and to each of the tissue interactomes. For each interactome, network embedding was computed by using the node2vec algorithm^59^. Each gene was sampled by 20 walks of length 10, and every interactome was represented by 64 vectors.

#### Expression variability features

Data of expression variability scores per gene and tissue were obtained from Simonovsky et al^60^. Scores reflected the variability in expression levels of a gene across samples of a given tissue, and were available for 19 tissues. Expression variability scores during development per gene and organ were computed based on Cardoso-Moreira et al^37^. Per organ and time point, each gene was assigned with its median normalized counts over the respective samples. Next, the expression variability of a gene in that organ was set to the coefficient of variation computed over the gene’s median normalized counts in all developmental time points of the given organ.

#### Paralogous genes’ features

Paralogous genes were defined as gene pairs whose reciprocal sequence identity was ≥ 40% according to Ensembl-BioMart. We assessed the quantitative relationships between paralogous genes as described in Barshir et al^42^. Specifically, we calculated per sample the expression ratio between a gene and its best-matching paralog, where the best-matching paralog was defined as the paralog with the highest sequence identity that was expressed in any tissue according to GTEx. Next, we created a feature per tissue where each gene was assigned with the median expression ratio computed across samples of that tissue. To account for genes with multiple paralogs, we calculated per sample the ratio between the expression of a gene and the summed expression of all its paralogs with over 40% reciprocal sequence identity. Next, we created another feature per tissue where each gene was assigned with the median ratio computed across samples of that tissue.

#### Differential process activity features

We associated genes with biological processes by using Gene Ontology^61^ (GO) terms and gene annotations, which were downloaded from Ensembl-BioMart. We favored specific rather than general biological processes and thus considered only GO terms to which 3-100 human genes were annotated. We associated each term with a differential activity score per tissue, which reflected the expression of its corresponding process in that tissue relative to other tissues. Per tissue, the differential activity score of a term was set to the average over the log2 fold-change value of its genes, where log2 fold-change value of each gene was computed based on its expression in that tissue relative to other tissues^62^. Using these scores, we associated each gene with the set of terms to which it was annotated, and created four gene features per tissue, as follows: (i) The minimum differential activity score of the gene’s terms in the given tissue. (ii) The maximum differential activity score of the gene’s terms in the given tissue. (iii) The median differential activity score of the gene’s terms in the given tissue. (iv) The mean differential activity score of the gene’s terms in the given tissue.

#### Data imputation

Imputation of missing values was achieved via a MICE-inspired iterative imputation function^63^. Each feature with missing values was considered as the target of a regression model, while using the rest of the features dataset as training data. First, per feature, all missing values were imputed by the median value of the respective feature. Then, per feature, the imputed values were reassessed by a Bayesian ridge regression model that used the 100 nearest features as training. Nearest features were evaluated based on the absolute correlation coefficient between each feature and the target feature. The reassessment of each feature was iterated 10 times, such that each iteration was initialized with values assessed in the preceding iteration.

#### Data transformation and scaling

Unlike tree-based models, logistic regression and neural network models could be confounded by nonsymmetrical distributions of data per feature. The main features that showed nonsymmetrical distributions were gene expression and preferential expression. These features were transformed to make them more symmetric by applying a Yeo-Johnson power transformation^64^. Transformation was applied after these data were used for the construction of other features and before applying the different machine learning (ML) models. Following transformation, each feature in the complete dataset was scaled per tissue to values between −1 and 1 while preserving the shape of its data distribution.

### The dataset of diseases, causal genes, and susceptible tissues

The set of Mendelian disease genes included genes with a germline aberrant form that is known to cause a disease and was obtained from OMIM^65^. Only Mendelian disease genes with a known molecular basis (OMIM Phenotype mapping key = 3) were included, resulting in 3,924 disease genes. The set of Mendelian diseases and their susceptible tissues (i.e., the tissue that clinically manifest the disease) was compiled from previously published manually-curated datasets ^42,28^. Next, we combined between the two sets, such that each disease gene that was causal for a disease with a known susceptible tissue was associated with that tissue, resulting 1,105 tissue-associated disease genes. To support the application of machine learning methods, we labeled genes per tissue *t* according to their causality for a disease that manifests in *t*: Genes that were causal for a disease that manifests in *t* where labeled as positive for *t*; all other genes were labeled as negative for *t* (Table S2). In the analyses per tissue model *t*, the set of non-causal genes included all genes except for Mendelian disease genes; The set of tissue non-causal genes included all Mendelian disease genes except for those that were labeled as positive for *t*.

#### The annotation of diseases to brain regions

Specifically for brain, we further curated brain disorders to brain regions. For this, we associated brain sub-regions that were sampled by GTEx with six distinct regions: Cortex (including anterior cingulate cortex (BA24), hippocampus, cortex, frontal cortex); cerebellum (including cerebellum, cerebellar hemisphere); basal ganglia (including caudate, nucleus accumbens, putamen); spinal cord; hypothalamus; and amygdala. Next, we manually associated brain disorders with their susceptible brain region(s). Associations were based on anatomical findings per disease that were detailed in disease pages of OMIM^66^. We assigned each association with a confidence level between 3 (high) and 1 (low), as follows: Associations based on clinical synopsis of the disease were assigned a confidence level of 3, unless they were described as pertinent only to some of the patients, in which case they are assigned a confidence level of 2. Associations based on disease description were assigned a confidence level of 2. Associations based on clinical features, which in some cases described finding relevant to a small subset of the patients, were assigned a confidence level of 1. Brain diseases that were not associated with any of the above regions (i.e., were associated with a different region, or were associated with brain but not with a specific region) were defined as ‘Other’. The union of all diseases that were associated with any region was designated as ‘whole brain’. The resulting dataset appears in Table S3.

To support the application of machine learning methods, we labeled genes per brain region *b* according to their causality for a disease that manifests in *b*: Genes that were causal for a disease that manifests in *b* at a confidence level of 2 and above were labeled as positive for *b*; all other genes were labeled as negative for *b* (Table S3).

### Using machine learning (ML) models to illuminate tissue-selectivity mechanisms

Below we describe the ML method used for interpretability analysis, its application to specific genes and to all tissue-causal genes, and the SHAP (SHapley Additive exPlanations) analysis of feature importance that was used to interpret the resulting models.

#### ML method for interpretability analysis

To create interpretable models we used the gradient boosted tree (GBT) algorithm. GBT trains a sequence of logistic regression trees, where each successive tree aims to predict the pseudo-residuals of the preceding trees assuming that the loss function is logistic loss. This method allows combining a huge number of shallow logistic regression trees by setting the learning rate to a small value. We employed the popular variant of GBT named ‘Extreme Gradient Boosting’ (XGBoost, XGB), which is considered the state-of-the-art algorithm for training GBT^67^. We applied XGB to interpret the tissue-selectivity of query genes, and the tissue-selectivity of tissue-causal genes for eight tissues, as described below. The input to XGB included the features dataset and the gene labels corresponding to the specific classification task. To reduce run time owing to the size of the features dataset and to reduce noise derived from non-contributing features, each application of XGB was preceded by feature selection that limited the number of relevant features per application to 50. Features were selected by applying support vector machines (SVM) with L1 regularization. We used SVM due to its capability to address high dimensional data. SVM was trained on, and fitted to, each relevant training set. By this, we selected different tailored sets of relevant features per application.

For more details on XGB implementation see ‘ML implementation details’ below.

#### XGB application to specific genes and to all tissue-causal genes

To interpret the tissue-selectivity of query genes (e.g., Fig 2), we trained an XGB model on all genes in our dataset except the query gene. For a query gene that is causal for a disease that manifests in tissue *t*, genes that were causal for diseases that manifests in *t* were labeled as positive, other genes were labeled as negative. The query gene was then tested by the model.

To interpret the tissue-selectivity of tissue-causal genes for a tissue *t* (e.g., Fig. 3A), we trained an XGB model on all genes in our features dataset. Genes that were causal for diseases that manifest in *t* were labeled as positive, all other genes were labeled as negative. We analyzed eight tissues for which the number of tissue-causal genes (positive genes) was sufficient for creating a robust model (i.e., over 60 tissue-causal genes). These tissues included blood, brain, heart, liver, nerve, skeletal muscle, skin and, testis. We tested the validity of the XGB model per tissue *t* by using 10-fold cross validation (Fig. S2).

#### SHAP analysis of feature importance

To identify the importance of the different features per model we applied SHAP tree algorithm to trained XGB models using default parameters^68^. To enable a summarized perspective on feature importance across different models, we normalized the SHAP feature values per model by dividing the value of each feature by the sum of values of all features in that model.

To identify recurrent patterns of features per model, we associated each feature with its feature type and with its tissue of origin. For example, the feature ‘brain preferential expression’ was associated with ‘preferential expression’ as its feature type and with ‘brain’ as its tissue of origin. Tissues-of-origin that were not part of the eight modeled tissues were considered as ‘Other’, and their corresponding features were grouped together. Next, to identify recurrent feature types per model, we summed up the normalized SHAP values of features belonging to the same feature type. Likewise, to identify recurrent tissues-of-origin per model, we summed up the normalized SHAP values of features belonging to the same tissue-of-origin.

### The TRACE ML framework for prioritizing tissue-selective disease-causing genes

Below we describe the TRACE framework and the application of TRACE to predict genes that are causal for tissue-specific diseases.

#### The TRACE framework

TRACE was composed of stacking of two layers. The first layer of TRACE consisted of five ML methods for training classifiers, denoted as base learners. The ML methods included logistic regression (LR), the tree-based ensemble methods XGB (described above), random forest (RF), and GBT (in the Scikit-learn implementation) initiated by a logistic regression model (LR+GB), and a multilayer perceptron (MLP) with one hidden layer. The second layer of TRACE consisted of a meta-learner MLP with two hidden layers.

In general, the input to TRACE included the features dataset and the gene labels corresponding to the specific classification task. Each base learner was applied independently to the input and produced a causality score per gene. Causality scores per base learner were scaled between 0 (non-causal) and 10 (causal). The output of the five base learners was the input to the meta-learner, which produced a final TRACE score per gene. The TRACE score was also scaled between 0 and 10.

To reduce run time owing to the size of the features dataset and to reduce noise derived from non-contributing features, all applications and folds of XGB, LR, and LR+GB were preceded by feature selection that limited the number of relevant features. Features were selected by applying SVM with L1 regularization. SVM was trained on and fitted to each relevant training set, which resulted in sets of relevant features that were tailored per application and per fold. To guarantee that the SVM-based feature selection will not entirely eliminate features that will contribute through non-linear relationships, feature selection was not applied to RF and MLP, which inherently handle non-contributing features by not selecting trees that rely on them (RF) or minimizing their weights (MLP).

#### TRACE application to genes that are causal for tissue-specific diseases

We applied TRACE to each tissue with over 60 tissue-causal genes, to predict the tissue-causality of genes. Per tissue *t*, the input to TRACE included the features dataset and the gene labels corresponding to their causality in that tissue: Genes that were causal for diseases that manifest in *t* were labeled as positive, all other genes were labeled as negative. We assessed the validity of each of the base-learner models and of the meta-learner model by using 10-fold cross validation (Fig. S3). Specifically, all genes in our dataset were randomly partitioned into 10 disjoint subsets, while preserving the ratio of tissue causal to tissue non-causal genes across subsets. Per fold, prior to running the base learners XGB, LR, and LR+GB, we performed feature selection, and then trained the models using the selected features. Per base-learner, the probabilities of genes within each subset to be tissue-causal in *t* were computed once by using a model that was trained on genes in the other nine subsets. Next, the scores of the five base-learners were used as features for the second level MLP meta-learner model. The meta-leaner model was trained on the scores of genes in the other nine subsets, and was then applied to predict the final TRACE scores of the genes within the given subset.

To evaluate each model we used the area under the receiver operating characteristic curve (AUC), where each point on the curve corresponded to a particular cutoff, representing a trade-off between sensitivity and specificity, and a summarized precision-recall score that computes the weighted mean of precisions achieved at each cutoff of the precision-recall curve.

#### Comparison to pBRIT^12^

We applied pBRIT to model each of the eight tissues that were modeled by TRACE. To mimic the input to TRACE per tissue, pBRIT was applied to the same distinct subsets that were used in TRACE. pBRIT was run via its web interface, by using the data fusion method of ‘TFIDF’. Query genes labels were withheld from the regression.

### ML Implementation details

All ML methods were implemented using the Scikit-learn python package^69^, except for XGB, which was implemented using the Scikit-learn API of the XGBoost package^67^. Per ML method, all hyper-parameters of the models were tuned manually to achieve higher AUC and summarized precision scores. Since AUC and summarized precision scores of the different tissue models showed small differences per method, tuning per method was done simultaneously for the eight tissue models. The same hyper-parameter values were then applied to all tissue models. For tissue-causality models and for patient-specific models, data contained mostly non-causal genes. To deal with class imbalance, the balance of positive to negative weights was set to 0.01 when training LR, XGB, RF, and the LR part of LR+GB.

#### XGB

The hyper-parameters of XGBoost were set to build a decision forest consisting of 150 trees. Each tree had a maximum tree depth of nine. Gamma was set to 0. To prevent overfitting, we set the step size shrinkage (eta) to 0.1.

#### RF

The number of trees was set to 1,000.

#### LR

LR was used with an lbfgs solver and maximum of 100,000 iterations.

#### LR+GB

LR+GB was implemented by using the ‘GradientBoostingClassifier’ function with LR parameters set with an lbfgs solver and maximum of 100,000 iterations and consisting of 80 trees.

#### MLP

The base-learner MLP was implemented by using the ‘MLPClassifier’ function with two hidden layers of size 10 each and ReLU activation function. Alpha was set to 0.5. Batch size was set to 200. The meta-learner MLP was similarly implemented except that alpha was set to 0.1 and the learning rate was adaptive and initiated at 0.01.

#### SVM

SVM was implemented by ‘LinearSVC’ function with a maximum of 10,000 iterations. For interpretability models, C was set to 2 and number of features was limited to 50. For TRACE prediction models C was set to 0.1.

### Analysis of tissue selectivity for distinct brain regions

We focused on the subset of diseases that were associated with brain regions with a confidence level ≥ 2, and on brain regions that were associated with at least 60 causal genes. TRACE analysis was similar to the analysis of other tissues. To assess the selectivity of diseases to a specific brain region, we compared between the scores of genes that were causal in the specific region (confidence level ≥ 2), to the scores of all other brain-causal genes (confidence level of 3), and to the scores of brain-causal genes that were not annotated to a specific brain region (denoted as ‘other’ in the manually-curated dataset, confidence level of 2).

### The application of TRACE to prioritize candidate causal genes in patients with rare tissue-selective diseases

#### Identification of candidate variants of unknown significance (VUS) per patient

We analyzed data from patients suffering from rare diseases that manifest in specific tissues. Patients were previously genetically diagnosed via exome sequencing and subsequent analysis as previously described^70–73^. The data per patient was de-identified, and variants were filtered as follows:

i. kept variants with call quality at least 20.0 in cases or at least 20.0 in controls AND outside top 5.0% most exonically variable 100base windows in healthy public genomes (1000 genomes).
ii. excluded variants that were observed with an allele frequency greater than or equal to 0.5% of the genomes in the 1000 genomes project OR greater than or equal to 0.5% of the NHLBI ESP exomes (All); or greater than or equal to 0.5% of the ExAC Frequency; or greater than or equal to 0.5% of the gnomAD Frequency; or filter variants unless established pathogenic common variant.
iii. kept variants (up to 20 bases into intron) that were experimentally observed to be associated with a phenotype: Pathogenic, possibly Pathogenic or disease-associated according to HGMD; or clinically relevant variants from CentoMD; or frameshift, in-frame indel, or stop codon change, or missense, or predicted deleterious by having CADD score > 15.0; or predicted to disrupt splicing by MaxEntScan; or within 2 bases into intron.
iv. In case of dominant genes, kept variants which are associated with gain of function, or hemizygous, or heterozygous, or heterozygous-amb, or compound heterozygous, or homozygous, or heterozygous-alt, or haploinsufficient and occur in at least one of the Case samples at the variant level; and not variants which are associated with gain of function, or hemizygous, or heterozygous, or heterozygous-amb, or compound heterozygous, or homozygous, or heterozygous-alt, or haploinsufficient, and occur in at least one of the control samples at the variant level in the Control samples.

In case of autosomal recessive genes, kept variants which are hemizygous, or compound heterozygous, or haploinsufficient, or homozygous, and occur in at least one of the case samples at the gene level in the Case samples; and not variants which are hemizygous, or compound heterozygous, or haploinsufficient, or homozygous, and occur in at least 1 of the control samples at the variant level in the Control samples.

Analyses were based on Ingenuity Variant Analysis version 5.4.20181019. Content versions: CADD (v1.3), Allele Frequency Community (2018-09-06), EVS (ESP6500SI-V2), Refseq Gene Model (2018-07-10), JASPAR (2013-11), Ingenuity Knowledge Base Snapshot Timestamp (2019-01-06 00:23:50.0), Vista Enhancer (2012-07), Clinical Trials (Stepford 190106.000), PolyPhen-2 (v2.2.2), 1000 Genome Frequency (phase3v5b), ExAC (0.3.1), iva (Oct 4 11:04 iva-1.0.736.jar), PhyloP (2009-11), DbSNP (151), TargetScan (6.2), GENCODE (Release 28), CentoMD (5.0), Ingenuity Knowledge Base (Stepford 190106.000), OMIM (May 26, 2017), gnomAD (2.0.1), BSIFT (2016-02-23), TCGA (2013-09-05), Clinvar (2018-08-01), DGV (2016-05-15), COSMIC (v86), HGMD (2018.3), SIFT4G (2016-02-23).

#### TRACE analysis of patient’s candidate genes

The data per patient was de-identified, and included the affected tissue of the patients and a list of VUS identified in that patient that remained after the filtering described in the preceding section. Each VUS was associated with its respective gene, denoted henceforth as candidate gene. Per patient, we created a TRACE model for each of its affected tissues (five of the 50 patients had two affected tissues). First, the patient`s candidate genes were entirely withheld from the features dataset, in order to be later prioritized by an independent TRACE model. The remaining, non-candidate genes, were labeled according to their causality in the patient’s affected tissue. In the first layer of TRACE, TRACE base learners predicted the tissue-causality of all non-candidate genes through 10-fold cross-validation procedure. Each base learner then predicted the tissue-causality of candidate genes by training a model on all non-candidate genes. In the second layer of TRACE, the scores of the base learners were used (once, no cross-validation), as features for the TRACE meta-learner. The meta learner was trained on all non-candidate genes, and was then used to predict the TRACE scores of the patient`s candidate genes.

#### Summary of TRACE results across patients

Per patient, we ranked patient’s candidate genes by their TRACE scores. Next, we associated the true causal gene of the patient with the percentile of its rank.

#### Comparison to GADO^13^

GADO prioritizes genes based on the phenotypes they cause, with phenotypes defined according to Human Phenotype Ontology (HPO) terms. For each patient we selected a HPO term that fitted with their phenotype and was supported by GADO, and then prioritized candidate genes for that term. Patients presenting multiple phenotypes were associated with one of the relevant HPO terms. Patients presenting a syndrome lacking a suitable HPO term were associated with a relevant HPO term based on literature search. Four cases were excluded from the analysis since their relevant phenotype was not supported by GADO.

### Statistical tests

To test the null hypothesis that TRACE scores of two distinct gene sets have similar probabilities to be smaller or greater than the other, we used the Mann-Whitney U test. Correction for multiple hypothesis testing was done via the Benjamini-Hochberg procedure. To test the null hypothesis that GADO^13^ ranking of the pathogenic gene was better or equal to TRACE ranking we used paired Wilcoxon test.

### TRACE webserver

The TRACE webserver was implemented in Python by using the Flask framework with data stored on a MySQL database. The website client was developed using the ReactJS framework and designed with Semantic-UI. The charts were displayed by the Google-Charts library. The TRACE webserver supports all major browsers. The webserver presents the TRACE scores of input genes in the user-selected tissue. TRACE scores were based on 10-fold cross validation and computed as described in section ‘*TRACE application to genes that are causal for tissue-specific diseases’*.

## ACKNOWLEDGEMENT

This study was funded by the Israel Science Foundation [317/19 to E.Y.-L.] and by a Ben-Gurion University grant [to E.Y.-L. and L.R.].

